# Binocular vs. monocular recovery experience differentially promote recovery from visual deficits in a mouse model of amblyopia

**DOI:** 10.1101/2021.02.24.432698

**Authors:** Jessy D. Martinez, Marcus J. Donnelly, Donald S. Popke, Daniel Torres, Sarah Sheskey, Brittany C. Clawson, Sha Jiang, Sara J. Aton

## Abstract

Altered visual experience during monocular deprivation (MD) profoundly changes in ocular dominance (OD) in the developing primary visual cortex (V1). MD-driven changes in OD are an experimental model of amblyopia, where early-life alterations in vision lead visual disruption in adulthood. Current treatments for amblyopia include patching of the dominant eye, and more recently-developed binocular therapies. However, the relative impact of monocular vs. binocular recovery experiences on recovery of function in V1 is not well understood. Using single-unit recording, we compared how binocular recovery [BR] or reverse occlusion [RO] of identical duration and content affects OD and visual response recovery in mouse binocular V1 after a period of MD. We also tested how BR and RO affected MD-driven alterations of parvalbumin expression, and visually-driven expression of cFos in parvalbumin-positive and negative neurons. Finally, we tested how BR and RO affected recovery of normal visual acuity for the two eyes in the context of visually-driven behavior. We find that BR is quantitatively superior with respect to normalization of V1 neurons’ OD, visually-driven cFos expression, and visual acuity for the two eyes. However, MD-driven changes in the firing rate and response properties of V1 principal neuron and fast-spiking interneuron populations do not recover fully after either BR or RO. Binocular matching of orientation preference also remains disrupted in V1 neurons after both forms of recovery experience. Thus BR and RO, analogs of differing treatment regimens for amblyopia, differentially impact various aspects of visual recovery in a mouse model for amblyopia.

**Significance Statement:** Amblyopia resulting from altered childhood eye function is a leading cause of lifelong vision loss. Treatment typically involves patching of the dominant eye (forcing monocular visual experience), and produces only partial recovery of vision. Using a well-established mouse model of amblyopia, we directly compared how two types of visual experiences influence recovery of visual function. Our findings suggest that binocular vs. monocular visual experience differentially effect restoration of normal visual responses in cortical neurons, visually-driven neuronal gene expression, and visual acuity. Understanding how the quality of recovery experience impacts visual system recovery in amblyopia should provide critical insights for clinical strategies for its treatment.

## Introduction

During critical periods of postnatal development, experience-driven synaptic plasticity shapes neural circuits in a manner that affects lifelong sensory and behavioral functions (1, 2). This has been extensively studied in primary visual cortex (V1) where during a critical period, brief monocular deprivation (MD; occlusion of one of the two eyes) shifts responsiveness of V1 neurons to favor the spared eye (3, 4). This phenomenon, known as ocular dominance plasticity (ODP), results from depression of deprived eye (DE) responses, followed by potentiation of spared eye (SE) responses (5, 6). These changes in responsiveness are associated with a transient decrease in cortical inhibition (7, 8). Closure of the critical period for ODP is thought to involve restoration of “mature” levels of cortical inhibition, which disrupts subsequent competitive plasticity of excitatory inputs (9–12).

ODP has served as a well-established model of the cortical changes associated with amblyopia, a visual disorder caused by an alteration in input to V1 from one of the two eyes during early childhood. Amblyopia can lead to long-term disruption of binocular vision and poorer visual acuity in adulthood, with limited treatment options (13–16). Standard treatments to promote recovery from amblyopia include dominant eye patching and – more recently - binocular visual experience, with varying levels of effectiveness reported for both (17–22). While animal models using MD have yielded mechanistic insights into the initial loss of visual function with amblyopia, an unanswered question is how visual experience impacts subsequent recovery of function in V1. For example, both reverse occlusion (RO; analogous to dominant eye patching) and binocular recovery (BR) promote varying levels of recovery of DE responsiveness in developing cat V1 (23–27). However, there is no consensus on which type of visual experience provides the most complete and rapid recovery.

To address this question, we aimed to directly compare the effects of BR and RO experiences - of matched duration and quality - on post-MD recovery of V1 visual response properties. The effects of critical period MD on plasticity in mouse V1 has been well described (4, 28, 29). Though the organization of V1 differs between mice and other species, features such as orientation tuning and OD are similar, and are similarly affected by MD (30–32). Following a 5-day period of MD, we conducted a side by side comparison of 5-day BR vs. RO, with similar amount and content of visual stimulation, on recovery of DE responses and other visual response properties. Using single-unit recordings in binocular V1 (bV1), we find that OD shifts caused by MD are reversed by BR, but not RO. MD leads to both depression of DE-driven firing rate responses, and enhancement of SE responses which is most evident in fast spiking (FS) interneurons. While DE response depression is not rescued by either BR or RO, BR reverses SE response enhancement in FS interneurons. Using immunohistochemistry for to quantify DE visually-driven cFos expression in bV1, we find MD-induced reductions in cFos in both pyramidal neurons and parvalbumin-expressing (PV+) interneurons after BR experience. Suppression of cFos induction by DE stimulation was partially reversed by BR (particularly among PV+ interneurons), but RO had no effect. PV expression itself was also decreased in bV1 after MD, suggesting plasticity in the FS interneuron population. BR did not reverse this decrease, but RO restored PV expression in bV1 to control levels. To assess the functional implications of these changes, we used behavioral optokinetic responses to measure visual acuity changes for the two eyes after MD, BR, and RO. As expected, we find that DE visual acuity is reduced after MD; DE acuity recovers after BR, but not RO. More surprisingly, SE visual acuity is significantly reduced after both BR and RO. These results demonstrate that monocular and binocular recovery experiences have striking differences with regard to outcomes in a mouse model of amblyopia, at the levels of both V1 physiology and behavior. These differential effects may be driven by the different impacts of BR and RO on the activity and function of PV+ FS interneurons in the bV1 network.

## Results

### MD-induced OD shifts are reversed by binocular, but not monocular, recovery experience

We first directly compared the effects of BR and RO on recovery of V1 neurons’ OD, following a 5-day period of MD. To induce a maximal OD shift, MD occurred over 5 days at the peak of the critical period for ODP in mice (P28-33, **Figure 1A**, left). To ensure similar quality and quantity of visual experience between BR and RO groups, these mice were placed for 4 h/day (starting at lights-on, i.e., ZT0-4) in a square chamber surrounded by LED monitors presenting high-contrast, phase-reversing oriented grating stimuli (8 orientations at 0.05 cycles/deg, reversing at 1 Hz). VE occurred daily over a 5-day recovery period (P33-38) for both BR and RO groups. To ensure a high proportion of wake time and to increase overall arousal (i.e. increasing the duration of visual stimulation) during this period of visual enrichment (VE; **Figure 1A**, right), the chamber itself contained a novel running wheel, manipulanda, and treats (31). After the 5-day recovery period, visual responses for stimuli presented to the right and left eyes were compared for neurons recorded in bV1 contralateral to DE occluded during initial MD. Consistent with previous reports (1, 5), 5-day MD induced a large OD shift in favor of the spared (ipsilateral) eye (**Figure 1B-D**). 5-day BR returned OD in bV1 to levels similar to age-matched Control mice with binocular vision, reversing the effects of MD. In contrast, OD distributions following RO were virtually unchanged from OD distributions after MD (**Figure 1B-D**).

**Figure 1.**
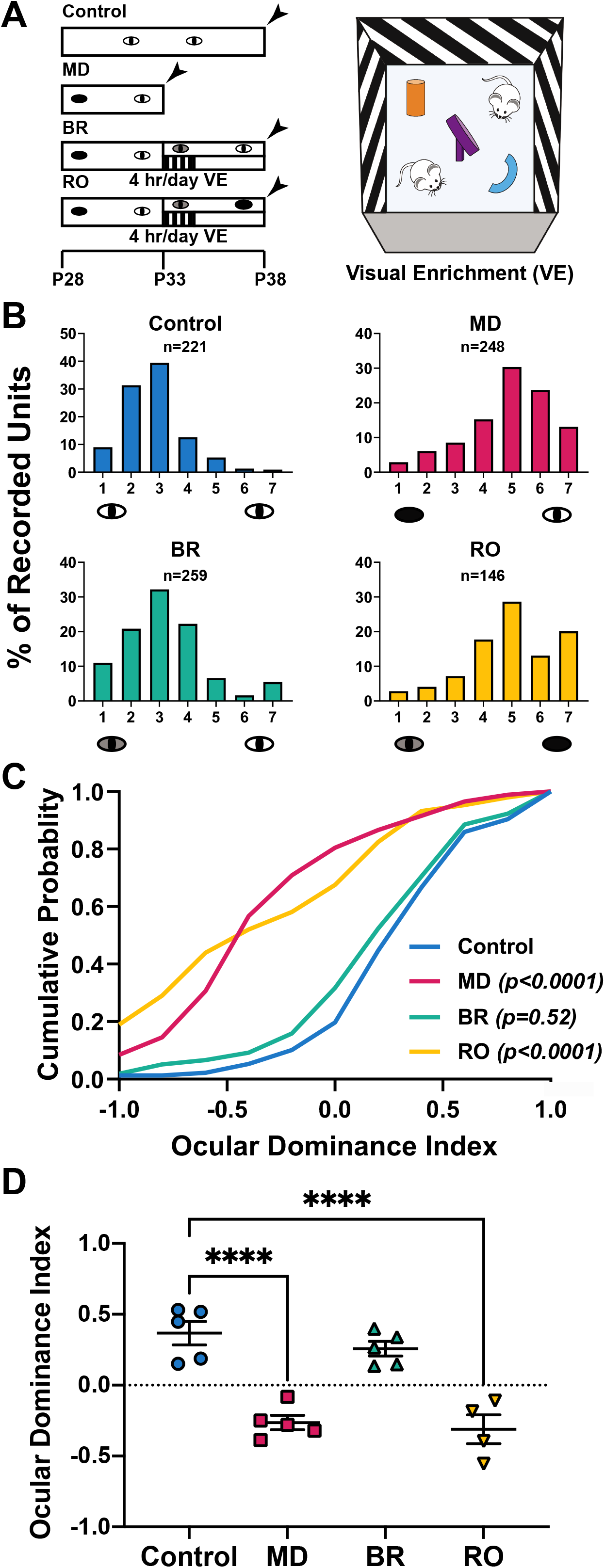
BR, but not RO, reverses OD shifts induced in bV1 after MD. ***(A, Left)*** Experimental design. Age-matched Control mice with normal, binocular vision were compared with mice that underwent 5-day MD from P28-P33. These mice were compared with two recovery groups with either binocular recovery (BR) or reverse occlusion (RO) experience. Recovery groups underwent daily 4-h periods of visual enrichment (VE), starting at lights on (ZT0) for 5 days after MD (P33-P38). Arrowheads indicate timing of either *in vivo* recording or visual acuity tests. ***(A, Right)*** VE consisted of visual stimulation with phase-reversing oriented gratings. Mice were given novel treats, toys, and a running wheel to encourage awake viewing of gratings during this period. ***(B)*** Ocular dominance (OD) histograms for single unit recordings from right-hemisphere bV1 (i.e., contralateral to the original DE) in all experimental groups. OD scores were distributed across a traditional seven-point scale, where 1 indicates neurons driven exclusively by the contralateral eye, 7 indicates neurons driven exclusively by the ipsilateral eye, and 4 indicates neurons with binocular responses. *n* = number of single neurons recorded in each group. ***(C)*** Cumulative graph of ocular dominance index (ODI) for all neurons shows OD shifts after 5-day MD (red line; *n* = 248 neurons from 5 mice) relative to Control mice with binocular vision (blue line; *n* = 221 neurons from 5 mice). BR mice (green line; *n* = 259 neurons from 5 mice) restores ODI to values comparable to Controls, while RO mice had ODI distributions similar MD mice (yellow line; *n* =136 units from 4 mice). *p* values indicate results of Kolmogorov–Smirnov test versus Control. ***(D)*** ODI per mouse averages for all groups. **** indicates *p* < 0.0001 vs. Control, Dunnett’s *post hoc* test; error bars indicate mean ± SEM.

### BR and RO differentially affect V1 principal neuron and fast-spiking (FS) interneuron firing rate responses

During initial shifts in OD, MD is known to effect a change in the balance of activity between principal neurons and FS interneurons (7, 8, 33). We compared how OD and visually-evoked firing in these cell populations change as a function of MD, BR, and RO. In our anesthetized recordings, FS interneurons (identifiable based on spike waveform features (7, 34)) represented roughly 20-25% of all stably-recorded neurons, across all treatment conditions (**Figure S1A**). We found that relative to neurons recorded from Control mice, MD led to significant OD shift in both principal neurons and FS interneurons. These changes, in both cell populations, were sustained in RO mice, and were reversed in BR mice (**Figure S1B**). Consistent with previous reports, MD also reduced the magnitude of DE-driven visually-evoked responses (5, 35); this change was present in both cell populations. DE firing rate responses remained depressed, and did not recover fully, after either BR or RO, for either cell population (**Figure 2A-B**; **Figure S2A-B)**. Spontaneous firing rates for principal neurons and FS interneurons (during presentation of a blank screen to the DE) showed a similar pattern – neurons recorded from MD, BR, and RO mice all showed depressed firing relative to those recorded from Control mice (**Figure S3A-B**; **Figure S4A-B**).

**Figure 2.**
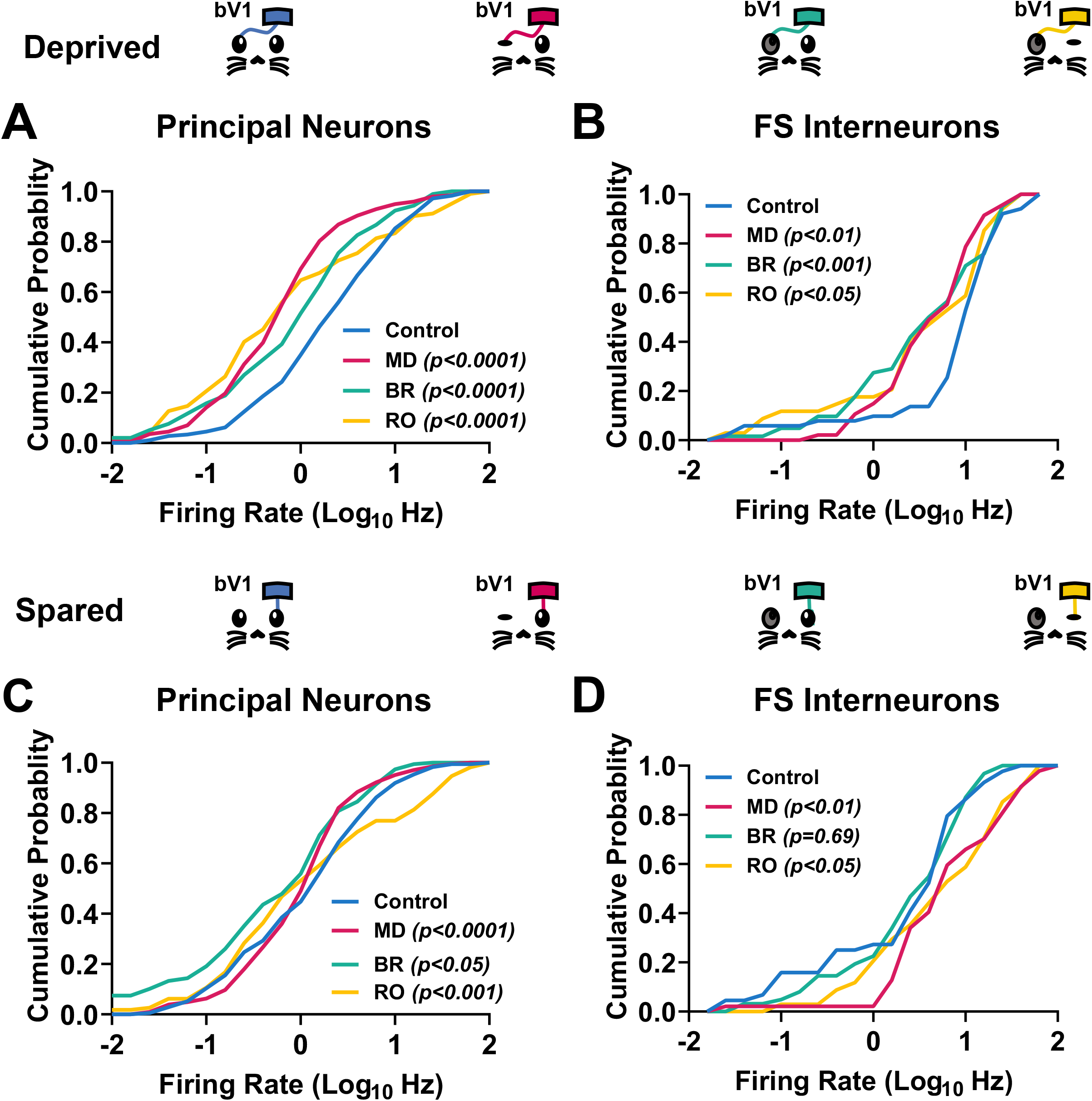
BR and RO differentially reverse MD-induced firing rate response changes in bV1 principal neurons and fast-spiking interneurons. ***(A)*** Cumulative distributions of maximal DE visually-evoked firing rates for principal neurons. Firing rates are significantly decreased after MD and decreases are maintained after both BR and RO. Sample sizes: Control (*n* = 170), MD (*n* = 201), BR (*n* = 197), RO (*n* = 102); *p* values indicate results of Kolmogorov–Smirnov test versus Control. ***(B)*** Cumulative distributions of maximal DE visually-evoked firing rates for FS interneurons. FS firing rates are significantly decreased after MD and these decreases are maintained after both BR and RO experience. Sample sizes: Control (*n* = 51), MD (*n* = 47), BR (*n* = 62), RO (*n* = 34); *p* values indicate results of Kolmogorov–Smirnov test versus Control. ***(C)*** Cumulative distributions of maximal SE visually-evoked firing rates for principal neurons, which were altered, relative to Control mice, in MD, BR, and RO groups. ***(D)*** Cumulative distributions of maximal SE visually-evoked firing rates for FS interneurons were increased, relative to Control mice, after MD. These changes persisted after RO, but were reversed by BR.

A consistent finding is that across species, prolonged MD results in response enhancement to SE stimulation in bV1 neurons (5–7, 35, 36). We found that SE response distributions in both principal neurons and FS interneurons were significantly altered after MD, with enhancement relative to Control mice most obvious in FS interneurons (**Figure 2C-D**; **Figure S2C-D**). After BR, principal neurons continued to show reduced SE firing rate responses among principal neurons, while firing rate responses in FS interneurons were renormalized to control levels. In contrast, in RO mice, firing rate changes in both cell populations caused by MD were sustained, with clear increases in firing responses in the FS population (**Figure 2C-D**; **Figure S2C-D**). Spontaneous firing rates (during presentation of a blank screen to the SE) showed a similar pattern. Principal neurons’ spontaneous firing was modestly affected in MD and RO mice only. In FS interneurons, significant increases in spontaneous firing were found after RO only (**Figure S3C-D**; **Figure S4C-D**).

To test how other aspects of visual responses were affected by MD and recovery experiences, we first calculated a metric of neuronal visual responsiveness (the responsiveness index; RI; **Figure S5**) for all recorded neurons. RIs (based on ratios between maximal and spontaneous firing rates) were generally increased for stimuli presented to the DE, and generally decreased for stimuli presented to the SE after MD. These changes were similar for both principal neurons and FS interneurons. RI changes for the DE were maintained in RO mice, although these changes were reversed (leading to decreased RIs for DE stimulation relative to Controls) in BR mice (**Figure S5A-B**). In comparison, SE RIs were generally reduced relative to Control mice, for both cell populations, in both RO and BR groups (**Figure S5C-D**).

### MD, BR, and RO differentially affect orientation tuning properties among V1 principal neurons and FS interneurons

Critical period plasticity associated with ODP also affects experience-driven changes in orientation tuning (35, 37). To test how orientation tuning in bV1 is differentially affected by recovery experiences, we next assessed orientation tuning changes in principal neurons and FS interneurons from each group. We found that MD selectively increased orientation selectivity (OSI90) for DE stimuli in the principal neuron population, but did not affect tuning in the FS population (**Figure S6A-B**). Consistent with previous reports (35), MD also significantly increased SE OSI90; this change was pronounced in both principal neurons and FS interneurons (**Figure S6C-D**). After BR, MD induced changes in OSI90 were generally reversed, for both eyes and in both neuron populations. After RO, OSI for DE stimuli was significantly increased in both populations relative to control mice (**Figure S6A-B**), and MD-induced OSI enhancements for SE stimuli were reversed (**Figure S6C-D**).

Consistent with recent reports (36, 38), we found that MD also reduced the degree of binocular matching of orientation preference (i.e., the matching of stimulus orientation causing maximal firing response between the two eyes) for bV1 neurons. This was true for comparisons of neurons with the most binocular response properties (i.e., those with OD values from −0.33 to 0.33; **Figure 3**), and comparisons across all bV1 neurons recorded (**Figure S7**). This aspect of V1 neuronal function did not recover completely in either BR or RO mice.

**Figure 3.**
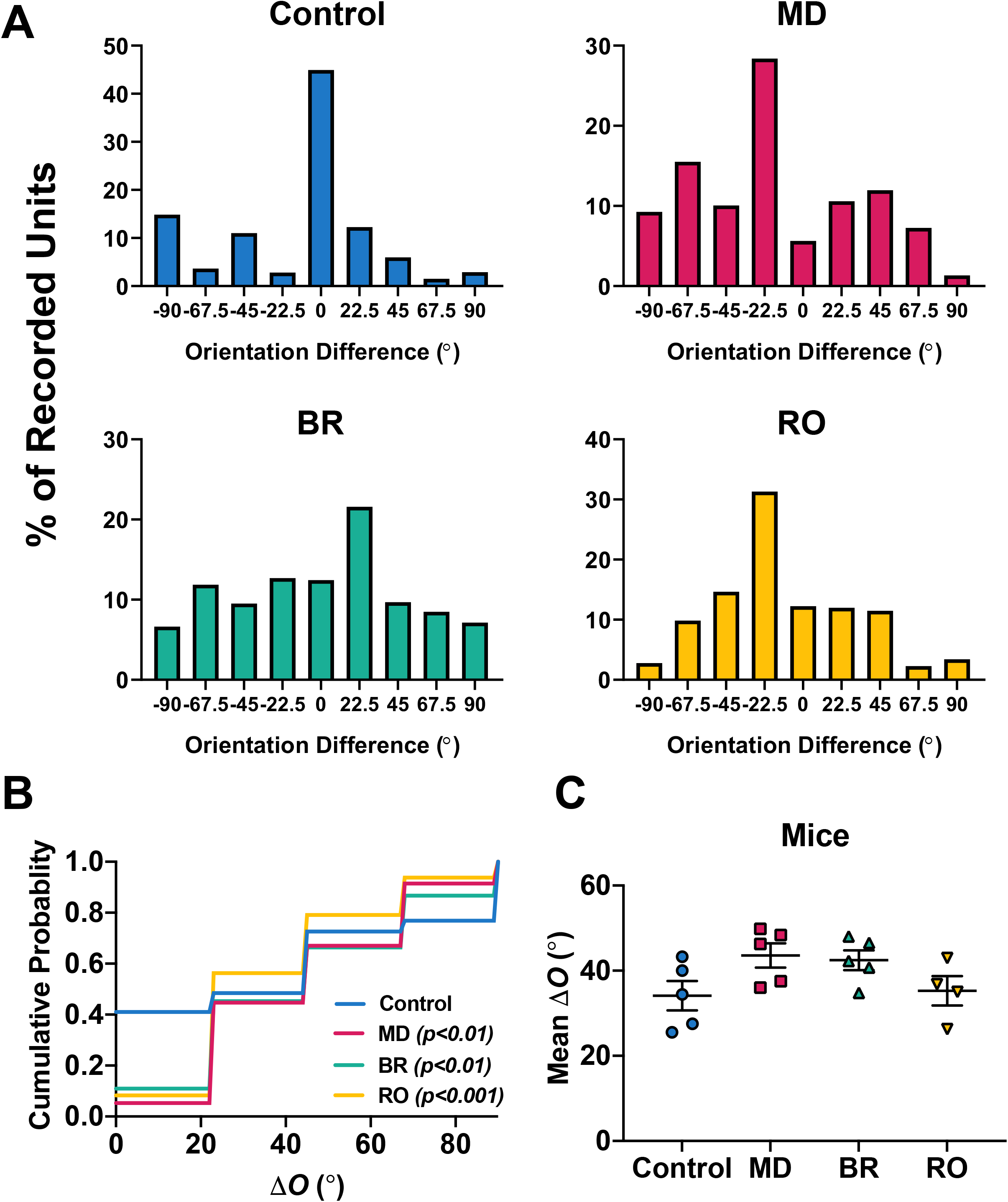
Neither BR nor RO restore binocular matching of orientation preference, which is disrupted by MD. ***(A)*** Plots showing distributions of differences in preferred orientation between the two eyes (in degrees), among neurons with relatively binocular responses (ODI values from −0.33 to 0.33) in all four groups. Control (*n* = 95 neurons, 5 mice); MD (*n* = 94 neurons, 5 mice); BR (*n* = 128 neurons, 5 mice); RO (*n* = 48 neurons, 4 mice). ***(B)*** Cumulative distributions of absolute values for binocular orientation preference differences (*Δ*O), showing increases after MD, BR, and RO relative to Controls. *p* values indicate results of Kolmogorov–Smirnov test versus Control. ***(C)*** *Δ*O per mouse averages for all groups. *F* = 2.6, *p* = 0.09, one-way ANOVA. Error bars represent mean ± SEM.

### Visually-driven cFos expression in bV1 PV+ interneurons is decreased after MD and recovered after binocular visual experience

Because our electrophysiological data suggested differential effects of BR and RO on the function of FS interneurons, we next examined the effects of MD and subsequent recovery experiences on activity-regulated protein expression in bV1 parvalbumin-expressing (PV+) interneurons. We used immunohistochemistry to quantify expression levels of both parvalbumin (which varies as a function of PV+ interneurons’ excitatory input (39)) and the activity-dependent marker cFos in bV1 (**Figure 4A**; **Figure S8**). Starting at lights on after a period of MD, BR, or RO, mice received 30 min of VE, with visual stimulation only to the DE. Mice were perfused 90 min later to characterize visually-driven expression of cFos. DE-driven cFos expression across bV1 was significantly reduced in bV1 after MD, consistent with previous reports (40, 41), and remained low after RO. In contrast, DE-driven cFos expression recovered at least partially after BR, with density of cFos+ neurons increasing above levels seen after MD or BR (**Figure 4A-B**). We also found that after MD, the density of PV-immunopositive neurons significantly decreased in bV1; this decrease was maintained after BR. However, PV-immunopositive cell density returned to levels seen in binocular vision Controls after RO (**Figure 4A,C**). Finally, to get a more direct measure of PV+ interneuron activation, we quantified the percentage of PV-immunopositive neurons which were cFos+. As expected from our electrophysiological data (**Figure 2**) MD significantly decreased DE-driven cFos expression in the PV+ population. While BR restored cFos expression in the PV+ population to levels seen in Control mice, DE-driven cFos expression reduced in RO mice (**Figure 4A,D**). Cell counts for cFos activated PV interneurons can be found in **Figure S8**.

**Figure 4.**
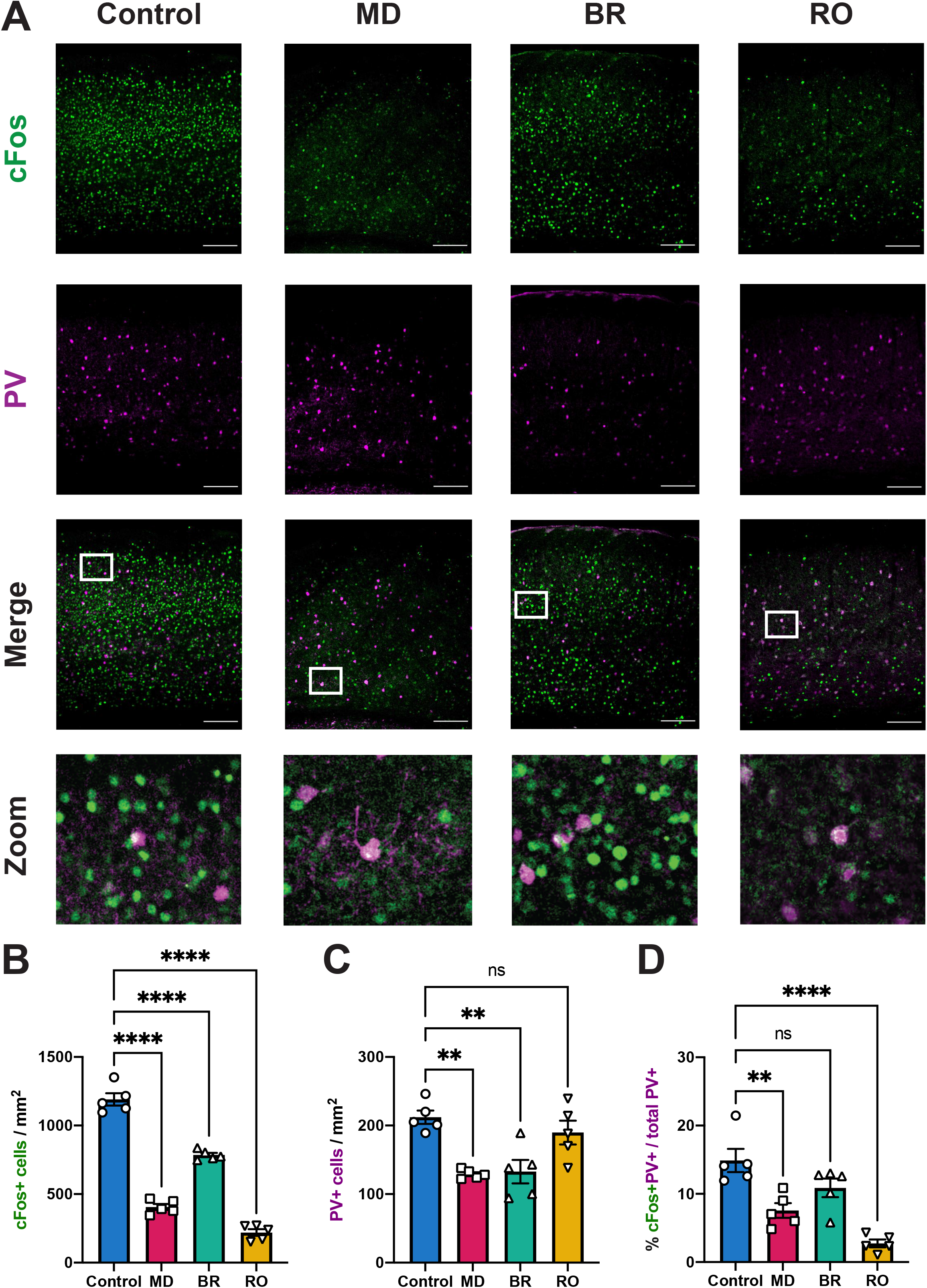
Parvalbumin expression and DE-driven cFos expression are reduced after MD and are partially restored after BR, but not RO. ***(A)*** Representative images of bV1 cFos and parvalbumin expression across treatment groups. Prior to sacrifice, mice had DE-only visual enrichment (see **Figure 1A**) for 30 mins, followed by 90 mins in their home cage. ***(B)*** DE-driven cFos+ neuron density is decreased in bV1 after MD. There is partial recovery of cFos expression after BR, but no increase after RO. ***(C)*** Density of bV1 interneurons with detectable parvalbumin expression decreases after MD. Decreases are maintained after BR, but not RO. ***(D)*** The percentage of parvabumin-expressing interneurons that are cFos+ after DE stimulation decreases after MD. These decreases are maintained after RO, but reversed by BR. Control (*n* = 5 mice); MD (*n* = 5 mice); BR (*n* = 5 mice); RO (*n* = 5 mice); error bars represent mean ± SEM. ** indicates *p* < 0.01, **** indicates *p* < 0.0001, Dunnett’s *post hoc* test

### Mice show differential recovery of visual acuity, measured behaviorally, with BR and RO

Because a common feature of amblyopia is long-lasting changes in visual acuity (42), we next tested how BR and RO differentially impacted visual acuity changes in mice after MD. We measured the visual acuity for each eye using an optokinetic behavioral paradigm. Vertical high-contrast sine wave gratings were presented in an interleaved manner to mice at different spatial frequencies, rotating in either a clockwise or counterclockwise direction **(Figure 5A)**, and optokinetic responses driven by the DE and SE were measured by an expert scorer blind to experimental conditions. As expected, 5-day MD reduced DE acuity, and had no significant effect on SE acuity, relative to Control mice. BR led to a partial recovery of DE acuity to Control levels, but significantly reduced SE acuity **(Figure 5B-C)**. In contrast, RO mice had reduced acuity for both eyes, relative to Controls **(Figure 5B-C).**

**Figure 5.**
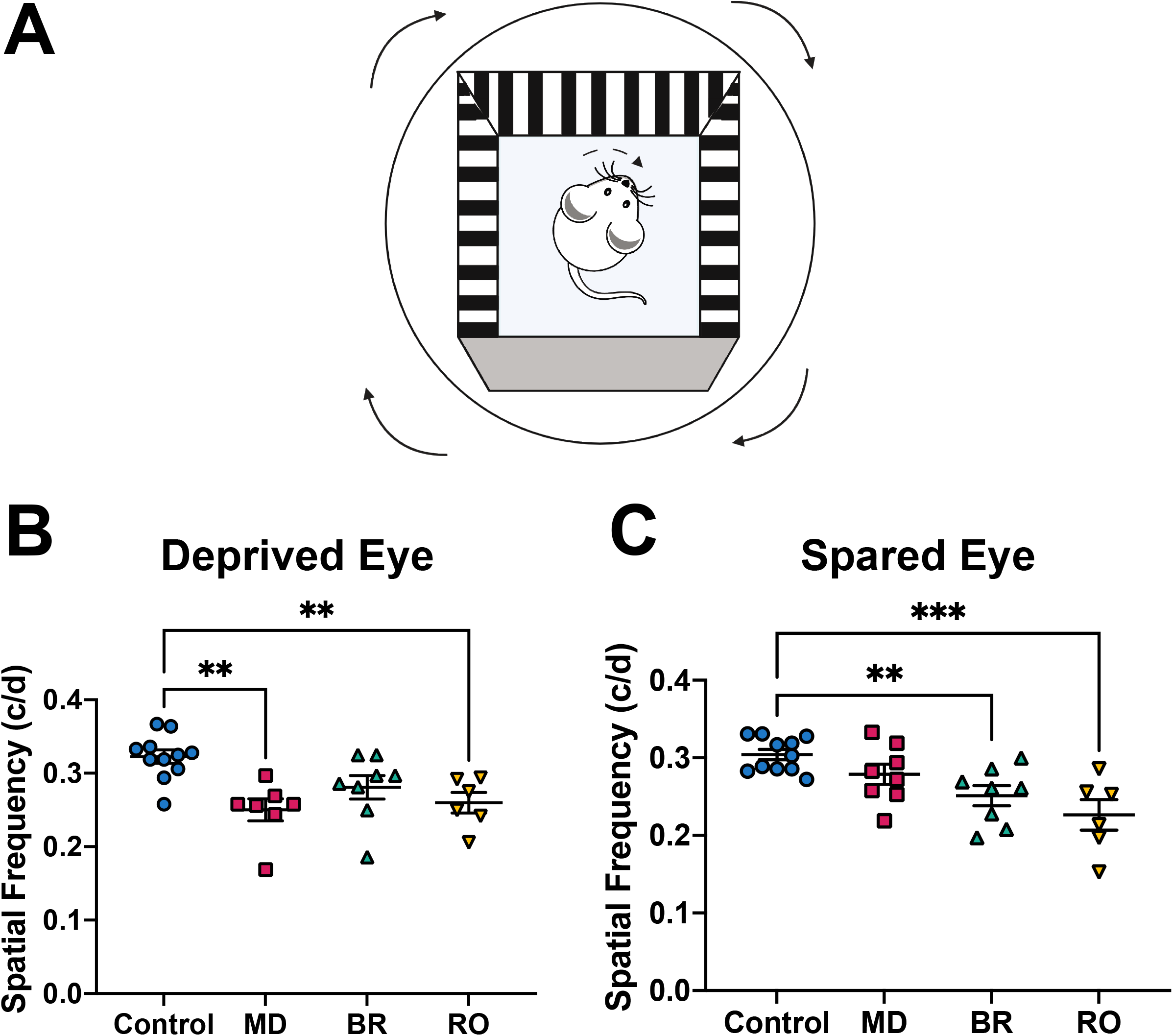
MD, BR, and RO differentially affect visual acuity for the two eyes. ***(A)*** Experimental setup. Optokinetic responses to clockwise and counterclockwise rotating gratings of different spatial frequencies were recorded from mice placed on an elevated platform in the center of the arena. ***(B)*** MD reduced DE acuity relative to Control mice. BR, but not RO, reversed MD-driven DE acuity decreases. ***(C)*** SE acuity for both BR and RO mice was reduced relative to Control mice. Control (*n* = 11 mice); MD (*n* = 8 mice); BR (*n* = 8 mice); RO (*n* = 6 mice).; Error bars represent mean ± SEM. ** indicates *p* < 0.01, indicates *p* < 0.001, Dunnett’s *post hoc* test

## Discussion

Here we made a side by side comparison of the effects of monocular (RO) vs. binocular (BR) visual experience on restoration of visual system function in a mouse model of amblyopia. Critically, we find that bV1 OD shifts in favor of the SE are reversed after BR, but are completely unaffected by RO **(Figure 1)**. These recovery OD shifts after BR experience are complete in principal neurons (with OD distributions statistically similar to Control mice after recovery), and at least partial in FS interneurons **(Figure S1)**. BR and RO also differentially affect firing rate response distributions and overall activity of principal neurons and FS interneurons. While BR experience partially reverses MD-initiated reductions in DE firing rate responses **(Figure 2)** and DE-driven cFos expression **(Figure 4)** in bV1, these effects of MD are (somewhat surprisingly) not reversed by monocular recovery experience (RO) using the previously deprived eye. MD-induced changes in SE-driven responses are also differentially affected by BR and RO. While increases in FS interneurons’ SE-driven firing rate responses after MD are not reversed by RO, they are reversed by BR **(Figure 2)**. However, many features of bV1 visual responsiveness remain disrupted after recovery using either method. For example, MD-initiated mismatching of orientation preference between the two eyes **(Figure 3)** and reduction in spontaneous firing rates **(Figure S3)** remain after both RO and BR. And ultimately, neither BR nor RO improves MD-initiated disruption of visual acuity for the two eyes **(Figure 5)**.

Because MD is known to initiate OD changes, in part, via gating of activity in the V1 PV+ interneuron network (7, 8), we aimed to test how both recovery experiences affected activity in PV+ interneurons vs. surrounding pyramidal neurons. We find that overall levels of parvalbumin expression across bV1 are reduced after MD, and that RO, but not BR, restores expression in PV+ interneurons to Control levels **(Figure 4)**. Previous work (e.g. in the rodent hippocampus) has demonstrated that parvalbumin expression levels vary cell by cell as a function of their excitatory input – with lower levels correlating with reduced excitatory drive (39). From this we conclude that RO leads to a resurgence in FS interneuron-mediated inhibition in the bV1 network, which is reduced after MD. BR, which is more permissive for OD shifts back to Control values, is associated with continued suppression of this inhibition. Consistent with the idea that BR suppresses neocortical inhibition (relative to RO mice), overall levels of cFos expression are significantly higher throughout bV1 in BR mice, vs. MD and RO mice **(Figure 4)**. In other words, resurgence of excitatory drive onto PV+ interneurons is correlated with no OD shift in RO, while continued reduction in PV+ interneuron activity is associated with restorative OD shifts in BR. This conclusion is consistent with recent data demonstrating that excitatory drive onto V1 PV+ interneurons acts as a gate which constrains ODP during, and outside of, the critical period (10, 43). Despite the restored parvalbumin expression levels we observe in RO mice (or possibly because FS+ interneurons have higher activity in bV1 after RO), DE-driven cFos expression is reduced – in both PV+ interneurons and surrounding pyramidal neurons – after RO. This may reflect both the continued suppression of DE responses after RO – consistent with the observed OD distributions in neurons recorded from RO mice – or also, altered levels of excitatory vs. inhibitory drive throughout the bV1 network.

Many factors can affect the degree of ODP initiated by MD in animal models of amblyopia, including behavioral state (35, 44, 45), and neuropharmacology (10, 46–48), and emerging data suggests that these factors may also affect recovery from amblyopia (47, 49, 50). However, recent findings have raised debate about whether dominant-eye patching, the decades-long standard of care for amblyopia, provides the optimal sensory stimulus for promoting recovery of vision (18, 22, 51, 52). Here, our side-by-side comparison of the effects of binocular vs. monocular visual experience in recovery of binocular visual responses and other aspects of vision. Our data show that while neither BR nor RO leads to complete restoration of all visual functions, the two forms of recovery experience have strikingly different effects on the function of bV1 and its responsiveness to visual input. They also suggest an underlying mechanism, by which RO and BR differentially engage the GABAergic circuitry within V1. We hope that these data will inform future studies for optimizing visual recovery treatment in children, which will lessen the long-term impact of amblyopia.

## Materials and Methods

### Animal housing and husbandry

For all experiments, C57BL/6 mice of both sexes were used. All mice were housed in a vivarium facility with a 12h:12h light/dark cycle and had *ad libitum* access to food and water. Mice were weaned at P21 and housed with at least one littermate of the same sex until surgical procedures or experiments. After surgeries and during recovery paradigms, mice were singly housed in standard cages with beneficial environmental enrichment. All mouse husbandry and experimental/surgical procedures were reviewed and approved by the University of Michigan Internal Animal Care and Use Committee.

### Monocular deprivation, recovery, and visual enrichment

For monocular deprivation (MD), mice (postnatal day 28) were anesthetized using 1-1.5% isoflurane. Nylon non-absorbable sutures were used to occlude the left eye. Sutures were checked twice daily to verify continuous MD. After MD (postnatal day 33), mice were anesthetized with 1-1.5% isoflurane a second time and left eyelid sutures were removed. Mice that underwent binocular recovery (BR) were then housed for 5 days with both eyes open. Mice that underwent reverse occlusion (RO) had the right (previously spared) eye sutured for a 5-day recovery period. Mice that lost sutures during the MD or recovery periods or developed eye abnormalities (e.g. cataracts) were excluded from the study. BR and RO mice underwent a similar 5-day period of enriched visual experience from postnatal day 33-38. This regimen consisted of a daily placement in a 15” × 15” Plexiglas chamber surrounded by 4 high-contrast LED monitors, from ZT0 (lights on) to ZT4. Phase-reversing oriented grating stimuli (0.5 cycles/deg, 100% contrast, 1 Hz reversal frequency) of 8 orientations were presented repeatedly on the 4 monitors in a random, interleaved fashion. During this 4-h period of daily visual stimulation, mice were encouraged to remain awake and explore the chamber via presentation of a variety of enrichment toys (novel objects, transparent tubes, and a running wheel) and palatable treats.

### *In vivo* neurophysiology and single unit analysis

Mice underwent stereotaxic, anesthetized recordings using a combination of 0.5-1.0% isoflurane and 1 mg/kg chlorprothixene (Sigma). A small craniotomy (1 mm in diameter) was made over right-hemisphere bV1 (i.e., contralateral to the original DE) using stereotaxic coordinates 2.6-3.0 mm lateral to lambda. Recordings of neuronal firing responses were made using a 2-shank, linear silicon probe (250 μm spacing between shanks, 32 electrodes/shank, 25 μm inter-electrode spacing; Cambridge Neurotech). The probe was slowly advanced into bV1 until stable, consistent spike waveforms were observed on multiple electrodes. Neural data acquisition using a 64-channel Omniplex recording system (Plexon) was carried out for individual mice across presentation of visual stimuli to each of the eyes, via a full field, high-contrast LED monitor positioned directly in front of the mouse. Recordings were made for the right and left eyes during randomly interleaved presentation of a series of phase-reversing oriented gratings (8 orientations + blank screen for quantifying spontaneous firing rates, reversing at 1 Hz, 0.05 cycles/degree, 100% contrast, 10 sec/stimulus). Spike data for individual neurons was discriminated offline using previously-described PCA and MANOVA analysis (53–56).

For each visually-responsive neuron, a number of response parameters were calculated (7, 35). An ocular dominance index (ODI) was calculated for each unit as (C-I)/(C+I) where C represents the maximal visually-evoked firing rate for preferred-orientation stimuli presented to the contralateral (left/deprived) eye and I represents the maximal firing rate for stimuli presented to the ipsilateral (right/spared) eye. ODI values range from −1 to +1, where negative values indicate an ipsilateral (SE) bias, positive values indicate a contralateral (DE) bias, and values close to 0 indicate similar responses for stimuli presented to either eye. A neuronal visual responsiveness index (RI) was calculated as 1-[(average firing rate at blank screen)/(average maximal firing rate at the neuron’s preferred orientation)]. An orientation selectivity index (OSI) was used to measure the sharpness of orientation tuning. OSI90 is calculated as 1-[(average firing rate at ± 90° from the preferred stimulus orientation)/(average maximal firing rate at the neuron’s preferred orientation)]. Finally, a measure of the difference in preferred orientation for stimuli presented to the two eyes (***Δ***O) was calculated, by subtracting the contralateral preferred orientation from the ipsilateral preferred orientation (values ranged from −90° to 90°). For quantitative comparison of these differences, we used the absolute values of these measures (i.e., 0°-90°)(38).

### Histology and immunohistochemistry

Following all electrophysiological recordings, mice were euthanized and perfused with ice cold PBS and 4% paraformaldehyde. Brains were dissected, post-fixed, cryoprotected in 30% sucrose solution, and frozen for sectioning. 50 μm coronal sections containing bV1 were stained with DAPI (Fluoromount-G; Southern Biotech) to verify electrode probe placement in binocular V1.

For quantification of parvalbumin and DE-driven cFos expression in bV1, mice from all groups underwent monocular suture of the original SE at ZT12 (lights off) the evening before visual stimulation. DE stimulation was carried out in the LED-monitor-surrounded arena with treats and toys to maintain a high level of arousal, as described for daily visual enrichment above. Mice from all groups were exposed to a 30-min period of phase-reversing oriented gratings as described above, after which they were returned to their home cage for 90 min (for maximal visually-driven cFos) expression prior to perfusion. Coronal sections of bV1 were immunostained using rabbit-anti-cFos (1:1000; Abcam, ab190289) and mouse-anti-PV (1:2000; Millipore, MAB1572) followed by secondary antibodies: Alexa Fluor 488 (1:200; Invitrogen, A11032) and Alexa Fluor 594 (1:200; Invitrogen, A11034). Stained sections were mounted using Prolong Gold antifade reagent (Invitrogen) and imaged using a Leica SP8 confocal microscope with a 10X objective to obtain images spanning the thickness of V1. Identical image acquisition settings (e.g. exposure times, frame average, pixel size) were used for all sections. cFos+ and PV+ cell bodies were quantified in 3-4 sections (spanning the anterior-posterior extent of bV1) per mouse using automated thresholding and cell counting scripts in ImageJ. Co-labeling was quantified using the Image J JACoP plugin (57). Quantification was based on average values across all sections imaged from each mouse.

### Optokinetic behavioral assay

Visual acuity was measured for the right and left eyes by measuring optokinetic responses using an OptoMotry tracking system (Cerebral Mechanics, Inc.) (58). For optokinetic measures, mice were placed on an elevated platform in the center of an enclosed arena surrounded by four LED monitors displaying a clockwise or counterclockwise drifting vertical sine wave gratings. Gratings were presented at multiple spatial frequencies and 100% contrast, in a randomly interleaved manner, and visual tracking behavior was measured by an expert scorer blind to each mouse’s experimental condition. Acuity was measured for each of the two eyes based on the spatial frequency threshold at which clockwise or counterclockwise tracking behavior ceased.

### Statistical analysis

Statistical analyses were carried out using GraphPad Prism software (Version 8.0). Comparisons of visual response properties were made for all stably recorded, visually-responsive units in bV1. Nonparametric tests were used for non-normal data distributions. Specific statistical tests and p-values can be found within the results section and in corresponding figures and figure legends.

## Data availability

All relevant data and analysis tools are available upon reasonable request from the corresponding author.

## Acknowledgments

We thank Abbey Roelofs (LSA Technology Services) for software programming assistance, and Chengmao Lin (Department of Ophthalmology) for coordination for experiments carried out in the Kellogg Eye Center Vision Research Core. This work was supported by a Rackham Merit Fellowship awarded to J.D.M., a Walt and Lilly Disney Award for Amblyopia Research from Research to Prevent Blindness to S.J.A., and R01 NS104776 awarded to S.J.A.

## Author Contributions

J.D.M and S.J.A. designed the research; J.D.M., M.J.D., D.S.P., D.T., S.S., B.C.C., and S.J. performed the research; J.D.M. and S.J.A. analyzed the data; J.D.M. and S.J.A. wrote the paper.

