## Supplemental Figures for "Binocular vs. monocular recovery experience differentially promote recovery from visual deficits in a mouse model of amblyopia"

**This PDF file includes:**

Supplementary Figure Legends

Figures S1 to S8

### Supplementary Figure Legends

**Figure S1: ODP measures for principal neurons and FS interneurons.** ***(A)*** Donut charts showing the proportion of recorded neurons classified as principal neurons and fast-spiking (FS) interneurons for all four groups. FS interneurons consistently make up roughly 20-25% of total recorded neuron population. Sample sizes (Principal neurons): Control ($n$ = 170); MD ($n$ = 201); BR ($n$ = 197); RO ($n$ = 102). Sample sizes (FS interneurons): Control ($n$ = 51); MD ($n$ = 47); BR ($n$ = 62); RO ($n$ = 34). ***(B)*** Ocular dominance index (ODI) values for principal neurons and FS interneuron populations. ***(Left)*** Principal neurons showed OD shifts in favor of the SE after MD, which were maintained after RO and are reversed by BR. ***(Right)*** OD shifts in FS interneurons after MD were maintained after RO, and were only partially reversed after BR. $p$ values indicate results of Kolmogorov–Smirnov test versus Control.

**Figure S2: Average DE and SE maximal firing rate responses in principal neuron and FS interneuron populations** ***(A-D)*** Per mouse average maximal firing rate responses (at the preferred stimulus orientation, for principal neurons and FS interneuron populations. DE maximal firing rate averages are significantly lower in MD, BR, and RO groups in the principal neuron population. No other significant differences were found at the level of per-mouse averages.  ** indicates $p < 0.01$, indicates $p < 0.001$, Dunnett's *post hoc* test

**Figure S3: Spontaneous firing rate changes in principal neurons and FS interneuron populations**. ***(A)*** Cumulative distributions of spontaneous firing rates for principal neurons, recorded during blank screen presentation to the DE. Firing rates were significantly decreased after MD and these decreases were maintained after both BR and RO experience. ***(B)*** FS interneuron DE spontaneous firing rates were significantly decreased after MD; these decreases were maintained after both BR and RO experience. ***(C)*** A significant decrease in principal neurons' SE spontaneous firing rates was found after MD, while a significant increase was found after RO. No change was seen after BR, relative to Control mice. ***(D)*** FS interneurons' SE spontaneous firing rates were significantly increased after RO experience. *p* values indicate results of Kolmogorov–Smirnov test versus Control.

**Figure S4: Spontaneous firing rate averages in principal and FS neuron populations.** ***(A-D)*** Per mouse average spontaneous firing rates for principal neurons and FS interneurons recorded in all conditions. Spontaneous firing during DE blank screen presentations was significantly lower after MD in principal neurons. No other significant differences were found at the level of per-mouse averages. ** indicates $p < 0.01$, Dunnett's *post hoc* test

**Figure S5: BR and RO differentially affect visual responsiveness (RI) in principal neurons and FS interneurons.** ***(A)*** DE RI values were significantly increased in principal neurons after MD; DE RIs remained increased after RO. BR decreases DE RIs relative to Control mice. *p* values indicate results of Kolmogorov–Smirnov test versus Control. Average values per mouse are shown below cumulative distributions of data. Error bars represent mean ± SEM. ***(B)*** DE RI values in FS interneurons were significantly increased after RO, and significantly decreased after BR, relative to Control mice. ***(C)*** SE RI values in principal neurons were significantly decreased after MD; these decreases remained after either BR or RO. ***(D)*** FS interneurons' SE RI values were significantly increased after MD, and significantly decreased after BR, relative to Control mice.

**Figure S6: BR and RO differentially affect orientation selectivity (OSI90) in principal neurons and FS interneurons.** ***(A)*** DE OSI90 values in principal neurons were significantly increased after MD and RO experience, and decreased after BR, relative to Control mice. *p* values indicate results of Kolmogorov–Smirnov test versus Control. Average values per mouse are shown below cumulative distributions of data. Error bars represent mean ± SEM. ***(B)*** FS interneurons' DE OSI90 values were unaffected by MD and BR, but were significantly increased after RO. ***(C)*** SE OSI90 values in principal neurons were significantly increased after MD; this change is reversed by both BR and RO. ***(D)*** FS Interneurons' SE OSI90 values were significantly increased after MD; this change was reversed by both BR and RO.

**Figure S7: Orientation preference matching for the two eyes across all recorded neurons**. ***(A)*** Plots showing distributions of differences in preferred orientation between the two eyes (in degrees), among all recorded neurons from all four groups. ***(B)*** Cumulative distributions of absolute values for binocular orientation preference differences ($\Delta$O), showing increases after MD, BR, and RO relative to Control mice. $p$ values indicate results of Kolmogorov–Smirnov test versus Control. ***(C)*** Mean $\Delta$O values, showing per mouse averages for all groups. $F = 2.2$, $p = 0.14$, one-way ANOVA. Error bars represent mean ± SEM.

**Figure S8: Raw count of cFos and parvalbumin-expressing interneurons in bV1 in each treatment group.** ***(A)*** Quantification of the number of double-labelled cFos+, parvabumin+ neurons in bV1 after DE visual stimulation. Control ($n = 5$ mice); MD ($n = 5$ mice); BR ($n = 5$ mice); RO ($n = 5$ mice). Error bars represent mean ± SEM; **** indicates $p < 0.0001$, Dunnett's *post hoc* test
